# Heart rate variability predicts decline in sensorimotor rhythm control

**DOI:** 10.1101/2021.01.08.424840

**Authors:** Marius Nann, David Haslacher, Annalisa Colucci, Bjoern Eskofier, Vinzenz von Tscharner, Surjo R. Soekadar

## Abstract

Voluntary control of sensorimotor rhythms (SMR, 8-12 Hz) can be used for brain-computer interface (BCI)-based operation of an assistive hand exoskeleton, e.g., in finger paralysis after stroke. To gain SMR control, stroke survivors are usually instructed to engage in motor imagery (MI) or to attempt moving the paralyzed fingers resulting in task- or event-related desynchronization (ERD) of SMR (SMR-ERD). However, as these tasks are cognitively demanding, especially for stroke survivors suffering from cognitive impairments, BCI control performance can deteriorate considerably over time. It would thus be important to identify biomarkers that predict decline in BCI control performance within an ongoing session in order to optimize the man-machine interaction scheme. Here we determine the link between BCI control performance over time and heart rate variability (HRV). Specifically, we investigated whether HRV can be used as a biomarker to predict decline during voluntary control of SMR-ERD across 17 healthy participants using Granger causality. SMR-ERD was visually displayed on a screen. Participants were instructed to engage in MI-based SMR-ERD control over two consecutive runs of 8.5 minutes each. During the second run, task difficulty was gradually increased. While control performance (*p* = .18) and HRV (*p* = .16) remained unchanged across participants during the first run, during the second run, both measures declined over time at high correlation (performance: -0.61%/10s, *p* = 0; HRV: -0.007ms/10s, *p* < .001). We found that HRV Granger-caused BCI control performance (*p* < .001) exhibited predictive characteristics of HRV on an individual participant level. These results suggest that HRV can predict decline in BCI performance paving the way for adaptive BCI control paradigms, e.g., to individualize and optimize assistive BCI systems in stroke.

## 1. Introduction

Brain-computer interface (BCI) systems allow for operation of hand exoskeletons to restore lost hand function, e.g., in finger paralysis after stroke (Ang et al., 2011). The best established method for such application utilizes voluntary control of sensorimotor-rhythms (SMR, 8-12 Hz) recorded by electroencephalography (EEG) (Birbaumer and Cohen, 2007). To gain control, stroke survivors are usually instructed to engage in motor imagery (MI) or to attempt moving the paralyzed fingers resulting in task- or event-related desynchronization (ERD) of SMR (SMR-ERD). Besides assistance, there is increasing evidence that regular use can trigger neural recovery leading to improved hand motor function (Ramos et al., 2012, Ang et al., 2015, Frolov et al., 2017a, Bai et al., 2020). However, optimal frequency, dose and intensity of BCI training to maximize the rehabilitation effect remains unclear (Young et al., 2015, Soekadar et al., 2015, Lang et al., 2016). Recently, it was proposed that merging both strategies, i.e., introducing assistance to perform bimanual activities of daily living (ADL) in the context of stroke rehabilitation protocols, may substantially increase the impact of BCI technology and broaden its use in the medical field (Soekadar, 2020, Soekadar and Nann, 2020, Soekadar, 2019)

However, BCI control tasks, e.g., performing repetitive MI of grasping movements, are cognitively demanding. As a consequence, decline in concentration and attention due to mental fatigue negatively affects BCI control performance (Myrden and Chau, 2015, Curran and Stokes, 2003). Especially stroke survivors suffering from cognitive impairments including post-stroke fatigue (Christensen et al., 2008, Acciarresi et al., 2014) have a limited capacity to maintain reliable BCI control over a longer period of time (Frolov et al., 2017b). Current research showed that subjective fatigue and inattention correlated with diminished BCI performance. This does not only reduce applicability of assistive BCIs after stroke but would also limit BCI training efficacy (Foong et al., 2019), as decline in attention was shown to negatively affect cortical plasticity (Stefan et al., 2004).

Despite extensive research efforts to identify predictive performance markers across subjects (Hammer et al., 2012, Blankertz et al., 2010), there is currently no adaptive control paradigm applied in BCI neurorehabilitation in which a critical within-session decline in SMR-based BCI performance becomes predicted on an individual level. Avoiding such performance decline would be crucial to sustain high control performance during assistance and to optimize rehabilitation efficacy. Maintaining optimal BCI control performance would be highly relevant for the management of the patient’s motivation to engage in regular use (Hammer et al., 2012) and to prevent overtraining in terms of dose and intensity (Soekadar et al., 2015). However, tracking negative trends in BCI performance over the course of a training session is difficult due to large trial-to-trial variations within each individual user (Grosse-Wentrup and Schölkopf, 2012). Moreover, the long-term goal is to apply BCI systems in everyday life environments, e.g., during activities of daily living (ADLs). Self-paced BCI control further impedes the detection of a negative trend as trial onsets are randomly initiated by the user and consequently more difficult to analyze. It would thus be important to identify sensitive biomarkers that predict decline in BCI performance at a point at which optimization of the patient-machine interaction scheme can yield sustained high control performance.

Recently, neuroadaptive human-computer interaction systems gained interest as they allow for *passive* cognitive monitoring without interfering with *active* voluntary control commands (Zander and Kothe, 2011, Gerjets et al., 2014). For instance, a neuroadaptive control paradigm has been developed to classify and adapt to the cognitive state of pilots during demanding flight simulations to increase their task accuracy (Klaproth et al., 2020). Neuroadaptive technology has been widely applied to discriminate between and adapt for different levels of workload during cognitive tasks, e.g., during a working memory task (Hogervorst et al., 2014). While cortical features are commonly used for classification, e.g., changes in the alpha (8-12 Hz) or theta (4-8 Hz) power band (Klimesch, 1999), also peripheral physiological measures as heart rate, heart rate variability (HRV), respiration rate or skin conductance were shown to be suitable for discrimination (Hogervorst et al., 2014).

Unlike EEG-based classification approaches commonly requiring a large number of electrodes and sophisticated analysis methods (Faller et al., 2019), peripheral physiological measures like heart rate or respiration rate are straightforward to record and less complex in analysis. Already over 150 years ago, Claude Bernard discovered the close interaction between the brain and the heart (Thayer and Lane, 2009). This functional link was found to be mediated by the vagus nerve originating from the brain and directly innervating the heart muscle. The vagus nerve has an important control function in slowing down pulse and heart excitability (countering the so called sympathetic innervation, which accelerates the pulse) to continuously adapt to physiological processes and account for changes in physical and/or cognitive demands. The strongest physiological adaptation process constitutes the adjustment of the heart rate to breathing, also called respiratory sinus arrhythmia (RSA). Since breathing induces unintended changes in blood pressure, the heart rate needs to be constantly adapted for pressure stabilization (Aasman et al., 1987).

To quantify the activity of the parasympathetic system, analysis of HRV is broadly applied. More precisely, there is strong evidence that particularly the high-frequency (HF-) HRV component in the range of 0.15 to 0.4 Hz reflects the modulation of vagus nerve activity (Berntson et al., 1997, Malik and Camm, 1993). In addition to physiological adaptation processes, numerous studies have shown that vagal activity is also sensitive to cognitive processes and attenuates during cognitively demanding tasks, e.g., during driving (Stuiver et al., 2014) and aviation simulation (Rowe et al., 1998, Muth et al., 2012), a visual attention test (Duschek et al., 2009) or a working memory and sustained perceptual attention task (Overbeek et al., 2014). A suppression of vagal activity induced by high cognitive demand results in lower adaptation to variations in blood pressure and thus less heart rate variability (Hogervorst et al., 2014).

However, despite extensive research in the field of psychophysiology and individual workload estimation based on HRV response patterns, it is unknown to what extend voluntary control of SMR in the context of BCI control can impact vagal activity. Moreover, it is unknown whether HRV can be used as a biomarker to predict decline in BCI control performance. Hence, understanding this relationship is a crucial prerequisite for the development of HRV-based neuroadaptive BCI systems to optimize control performance and stroke neurorehabilitation protocols. Here, we tested in a MI-neurofeedback study with 17 healthy participants whether decrease in HF-HRV predicts a decline in BCI control performance within the same session. The first objective (O1) was to evaluate specificity of HRV decline for BCI control performance. Second, we tested whether HF-HRV predicts average BCI control performance on a participant level within a single run consisting of 50 BCI control trials (O2). For this, we performed a Granger causality analysis to investigate the predictive characteristics of HRV. Granger causality, introduced by Clive Granger in 1969 (Granger, 1969), is a statistical hypothesis test and has been widely applied in economic science as a forecasting method, e.g., of two stock market time series. In short, a variable X *Granger-causes* a second variable Y, when prediction of Y is better including information of X than without. This is done by a series of F-tests on lagged values. Recently, Granger causality has gained broad interest and various applications in the field of neuroscience (Seth et al., 2015).

## 2. Methods

### 2.1 Participants

24 able-bodied participants (seven females, mean age 33±12 years) were invited to the University Hospital of Tübingen, Germany, for a familiarization and experimental session on two consecutive days. During the familiarization session, all participants gave written informed consent and were pre-screened whether they were able to elicit detectable SMR-ERD and whether they showed a normal heartbeat pattern to allow for HRV analysis. In case of medication intake on a regular basis or neurological diseases, participants were excluded from the study. After successful pre-screening, 17 participants were invited to the experimental session on the following day. The study protocol was in line with the Declaration of Helsinki and was approved by the local ethics committee at the University of Tübingen (registration code of ethical approval: 201/2018BO1).

### 2.2 Task description and study design

The participants were instructed to engage in motor imagery (MI) of left or right grasping movements and received visual feedback based on SMR-ERD modulations presented on a screen in front of them. Participants performed two consecutive runs with 50 trials each resulting in an overall duration of 500 s (8.5 minutes) per run. A run consisted of a pseudorandomized and externally paced sequence of MI and relax tasks with participants instructed to blank their minds and rest. Both tasks had a length of five seconds and were separated by intertrial intervals (ITI) with a randomized duration of 4-6 s.

The study followed a within-participant repeated measures one-factor design with task difficulty as independent variable. To increase task difficulty between runs, SMR-ERD detection threshold as well as ratio in number of MI to relax tasks were manipulated. During the first run, SMR-ERD detection threshold was kept constant and task ratio was balanced (task difficulty: ‘normal’). To increase task difficulty during the second run (task difficulty: ‘hard’), SMR-ERD detection threshold was gradually lowered by −20 % relative to the individual SMR-ERD detection threshold determined during calibration every 100 s. To avoid distortion of true negative rate, detection threshold for the relax tasks was kept unchanged. Moreover, task ratio was unbalanced with 60 % MI to 40 % relax tasks (overview of study design see Figure 1). The increase in task difficulty during the second run was not communicated to the participants to avoid biasing their motivational state or influence with their expectations.

**Figure 1:**
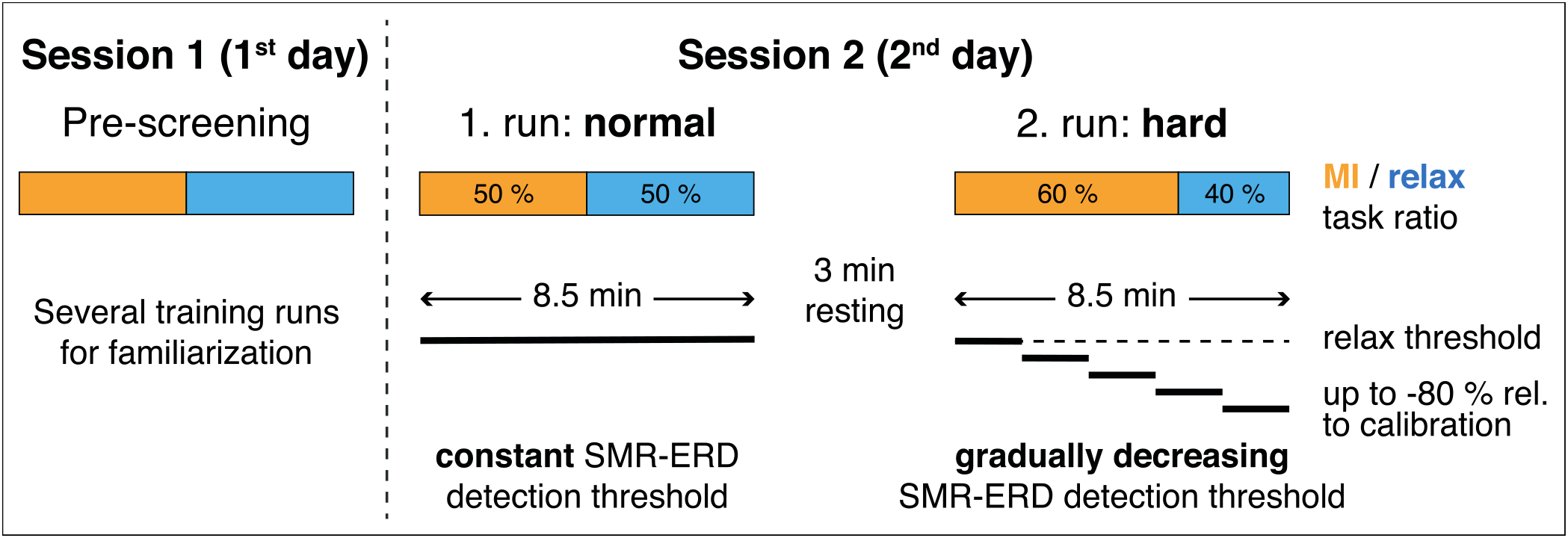
Overview of study design. Participants were invited for a pre-screening session (session 1) followed by an experimental session (session 2) on two consecutive days. During the pre-screening session, participants were familiarized with the setup and the motor imagery (MI) neurofeedback task. On the following day, participants performed two 8.5 min (500 s) long runs with different task difficulty; normal and hard. Task difficulty was increased by altering the ratio of MI to relax tasks (60/40 %) and by gradually decreasing the sensorimotor rhythms-SMR-) event-related desynchronization (ERD) detection threshold every 100 s by −20 % up to −80 % relative to the individual SMR-ERD detection threshold determined during calibration. Relax threshold was not altered (horizontal dashed line).

### 2.3 Brain-computer interface (BCI) control

Electroencephalography (EEG) was recorded from 22 conventional recording sites (Fp1, Fp2, AFz, F3, Fz, F4, T7, C3, Cz, C4, T8, CPz, P7, P3, Pz, P4, P8, POz, O1, O2, M1, M2 according to the international 10/20 system) and referenced to FCz. Ground electrode was placed at Fpz. Dependent on the selected laterality of the imagined grasping movements, either C3 for the right hand or C4 for the left hand together with the respective four nearest electrodes for Laplacian filtering were selected for further signal processing (McFarland, 2015). Two additional channels were used to assess the heart rate for offline HRV analysis (see section 2.4). All biosignals were amplified with a wireless saline-based EEG system (smarting®, mBrainTrain®, Belgrade, Serbia, in combination with a customized GT Gelfree-S3 EEG cap, Wuhan Greentek Pty.Ltd., Wuhan, China) and sampled at 500 Hz. All impedances were kept below 10 kΩ for high signal quality. To provide visual online feedback of participant’s voluntary SMR control, the BCI2000 software platform was used for real time processing and classification (Schalk et al., 2004).

First, EEG signals at C3 or C4 (dependent on the imagined hand) were bandpass-filtered at 1 to 30 Hz as well as Laplace-filtered to increase signal-to-noise ratio (McFarland, 2015, Soekadar et al., 2016). Afterwards, instantaneous SMR power amplitude of a 3 Hz-wide bin around the frequency of interest (FOI ± 1.5 Hz) was estimated from a 400-millisecond moving window based on an autoregressive model of order 50 according to the Burg algorithm (Soekadar et al., 2011). FOI was individually determined during calibration and was set to the frequency showing the largest SMR power downmodulation during MI. Lastly, SMR-ERD was calculated according to the power method of Pfurtscheller and Aranibar (1979).

BCI online control required individual calibration before the two experimental runs. Therefore, participants performed three short calibration runs, in which they followed 20 externally paced visual cues instructing them to either engage in MI or to relax. Based on the first run, in which no online feedback was provided, individual average power was estimated to normalize instantaneous power for SMR-ERD computation. During the following two calibration runs, an individual SMR-ERD detection threshold was determined to provide meaningful online feedback. Detection threshold was set to the average of elicited SMR-ERD across all MI tasks (Nann et al., 2020, Soekadar et al., 2016).

### 2.4 Analysis of heart rate variability (HRV)

For the analysis of heart rate variability (HRV), a one-lead electrocardiography (ECG) was derived with disposable Ag-AgCl electrodes (COVIDIEN plc, Dublin, Ireland) from the outer clavicles according to Mason and Likar (1966). ECG was lowpass-filtered at 130 Hz including removal of 50 Hz line noise. Interbeat intervals (IBIs) were extracted and corrected, in case IBIs exceeded the average IBI value of a window comprising 20 previous and 20 following beats by 30 % (von Tscharner and Zandiyeh, 2017). Cleaned IBIs were resampled at 4 Hz by using a modified Akima cubic interpolation to obtain equidistant time series (Berntson et al., 1997, Camm et al., 1996, Singh et al., 2004). The resampled IBI time series was bandpass-filtered at 0.15 to 0.4 Hz resulting in a time series that only contained the high-frequency (HF-) HRV component, which was found to reflect modulation in vagal tone (Malik and Camm, 1993). To receive the final HRV outcome measure, standard deviation of the filtered IBI time series was computed (Shaffer and Ginsberg, 2017). According to recent recommendations, no correction for respiration rate was performed since spontaneous beathing was present throughout the entire experiment (Laborde et al., 2017, Thayer et al., 2011). Since interindividual distribution of HRV measures are usually strongly positively skewed, HRV values are ln-transformed to regain normal distribution for further parametric statistical analysis (Riniolo and Porges, 2000).

### 2.5 Outcome measures

To capture temporal HRV changes within each experimental run (run length equals 500 s), HRV measures were computed based on overlapping 100-seconds windows iteratively shifted by ten seconds. This results in a HRV time series with 41 data points per run. Estimating HRV measures on the basis of such relatively short observation windows recently gained interest as it allows to study HRV changes with a higher temporal resolution compared to standard window lengths (Shaffer and Ginsberg, 2017). Several studies showed that such ultra-short-term HRV measures highly correlate with the clinically accepted short-term measures computed over a standard window length of five minutes (Munoz et al., 2015, Schippers et al., 2018, Castaldo et al., 2019).

The outcome measure for BCI performance was defined as the percentage of time, the SMR-ERD detection threshold was exceeded during the MI task and not exceeded during the relax task relative to trial length. To obtain the equal 100-second window length as for HRV data analysis, BCI performance measures were averaged across ten tasks (on average a task and a ITI resulted in ten seconds duration). Shifting this window iteratively by ten seconds results in a time series of 41 data points per run, equal to the length of the HRV time series.

For multilevel model analyses (see section 2.6) and plotting, both outcome measures were expressed as difference scores where values are subtracted from the first computed value of each run (representing difference to first 100 s) (Overbeek et al., 2014). Looking at difference scores instead of absolute values reveals changes within each run and facilitates comparison between levels of task difficulty.

### 2.6 Offline analysis

To investigate whether BCI performance as well as HRV changed over time and task difficulty, two separate multilevel models for each outcome measure were applied (Snijders and Bosker, 1999, Raudenbush and Bryk, 2002) using the package ‘nlme’ in R (Pinheiro et al., 2013). Multilevel modeling is an extension of the linear regression analysis and constitutes a powerful and flexible method to analyze hierarchical and repeated-measures data (Muth et al., 2012). In our study, the dependent variables ‘BCI performance’ and ‘HRV’ as outcome measures (level 1) were nested within ‘participants’ and ‘task difficulty’ residing at level 2. In addition, the predictor variables ‘time’ (41 continuous data points) and ‘task difficulty’ (two categorical levels: normal, hard) resided at level 1. Multilevel modeling can account for the lack of independence of the outcome measures within participants and is thus well suitable for a repeated-measures design. Additionally, unlike a repeated-measures ANOVA, the assumption of sphericity can be neglected. To account for the high autocorrelation of both outcome time series, which was caused by the moving windows, a first-order autoregressive covariance structure was modeled (Field et al., 2012). Normality of residuals was checked by visual inspection with quantile-quantile plots (Q-Q plots) and scatter plots of fits vs. residuals. To test for an overall effect of predictor variables onto the response variable, the model was compared with the baseline model (i.e., no predictors at level 1 or 2) using a chi-square likelihood ratio test.

To test whether HRV predicted BCI control performance, Granger causality analysis was applied (Granger, 1969). It was tested for whether the null hypothesis H_0_, which states that HRV does not *Granger-cause* BCI control performance, could be rejected. However, Granger causality analysis can indicate spurious causality when one or both time series are non-stationary (He and Maekawa, 2001). Since both outcome measures were assumed to change over time, i.e., they show a trend, non-stationarity of time series was presumed. Toda and Yamamoto (1995) introduced a method, which allows to test for Granger causality on non-stationary time series. This method is based on augmented vector autoregression (VAR) modeling and a modified Wald test statistic and was shown to be superior to the original Ganger test (R packages ‘vars’ and ‘aod’). The Toda-Yamamoto implementation to test for Granger causality was applied on time series on single participant level. To combine test statistics of all participants, Fisher’s combined probability test (Fisher’s method) was applied to perform a meta-analysis of *p*-values (Fisher, 1925) (R package ‘metap’). Significance threshold for all statistical tests was assumed at *p* = .05.

## 3. Results

### 3.1 Changes of BCI performance over time

The comparison with the baseline model revealed a significant effect of ‘time’ and ‘task difficulty’ on ‘BCI performance’ (χ^2^(2) = 401.60, *p* < .0001). While there was no overall main effect of ‘time’ on ‘BCI performance’ (*b* = −0.087 [95 % CI: −0.202, 0.029], *t*(1324) = −1.47, *p* = 0.142), there was a significant interaction of ‘time’ and ‘task difficulty’ (*b* = −0.523 [95 % CI: −0.686, −0.359], *t*(1324) = −6.27, *p* = 0), indicating that ‘task difficulty’ affects the relationship between ‘time’ and ‘BCI performance’. To investigate the effect over ‘time’ for each run individually, interaction was resolved by conducting separate multilevel models for *normal* and *hard* ‘task difficulty’. The models were identical to the main model but excluded the main effect and interaction term of ‘task difficulty’. These analyses showed that during the *normal* run, there was no change in BCI performance over time (*b* = −0.087 [95 % CI: −0.212, 0.039], *t*(662) = −1.36, *p* = .176). However, during the *hard* run, there was a strong decline in BCI performance over time (*b* = −0.614 [95 % CI: −0.723, −0.504], *t*(662) = -10.97, *p* = 0). The gradient of the regression line implies an average decline in BCI performance of −0.6 % per 10-seconds, which accumulates to an overall drop of −24 % at the end of the *hard* run (compared to a non-significant decrease of −3.5 % in total during the *normal* run, Figure 2).

**Figure 2:**
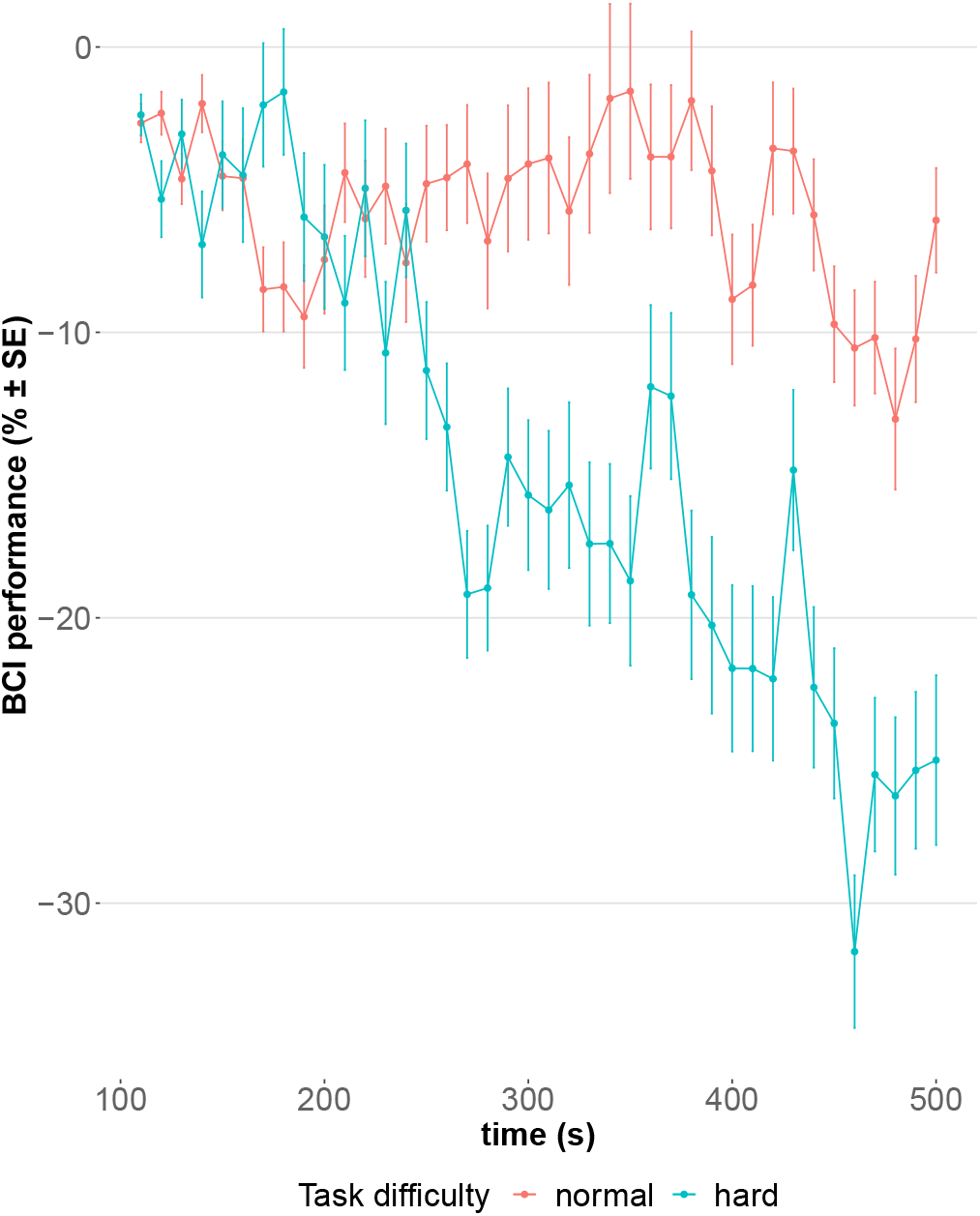
Change in brain computer interface (BCI) performance over time during runs with normal (red) and hard (blue) task difficulty level. Every run had a length of 500 s. Each point represents an average value (with error bars showing standard error, SE) across participants based on a 100-seconds window. Time series was generated by moving windows iteratively shifted by 10-seconds. First value of time series was computed based on the first 100 s and utilized as reference value for difference score calculation (i.e., subsequent values are subtracted from the first computed value).

### 3.2 Changes of HRV over time

Equal to the analysis of BCI performance, there was a significant effect of ‘time’ and ‘task difficulty’ on ‘HRV’ (χ^2^(2) = 94.71, *p* < .0001), but no overall main effect of ‘time’ was found (*b* = −0.002 [95 % CI: −0.006, 0.001], *t*(1324) = −1.30, *p* = 0.193). Even though not significant, there was a trend for ‘task difficulty’ affecting the relationship between ‘time’ and ‘HRV’ (*b* = −0.004 [95 % CI: −0.009, 0.001], *t*(1324) = −1.76, *p* = .079). Therefore, interaction was again resolved to investigate effects over time for both levels of ‘task difficulty’. These analyses revealed that during the *normal* run, there was no change in HRV (*b* = −0.002 [95 % CI: −0.005, 0.001], *t*(662) = −1.42, *p* = 0.157). During the *hard* run, however, HRV declined over time (*b* = −0.007 [95 % CI: −0.01, −0.003], *t*(662) = −3.40, *p* < .001, Figure 3). Thus, HRV shows the same behavior over time as BCI performance indicating that decline in HRV is specific to a decline in BCI performance.

**Figure 3:**
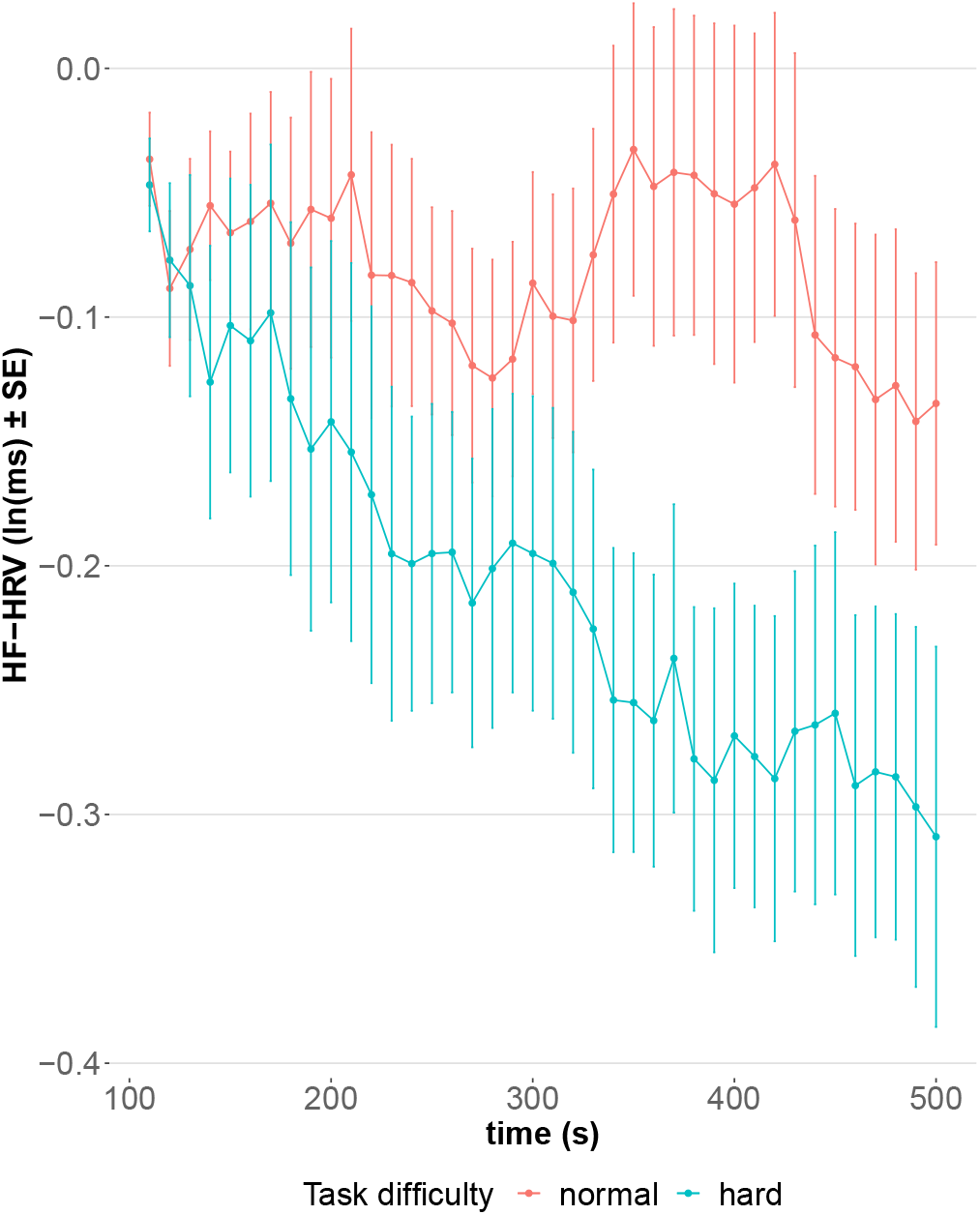
Change in natural logarithm (ln)-transformed high-frequency (HF) heart rate variability (HRV) over time during runs with normal (red) and hard (blue) task difficulty level. Every run had a length of 500 s. Each point represents an average value (with error bars showing standard error, SE) across participants based on a 100-seconds window. Time series was generated by moving windows iteratively shifted by 10-seconds. First value of time series was computed based on the first 100 s and utilized as reference value for difference score calculation (i.e., subsequent values are subtracted from the first computed value).

### 3.3 Time series prediction

Test statistic based on the Fisher’s method showed that HRV *Granger-caused* BCI control performance considering all participants including both levels of task difficulty (χ^2^(68) = 997.9, *p* < .0001). All *p*-values except one were highly significant according to the Toda-Yamamoto implementation. Figure 4 shows the time series of HRV and BCI performance of one participant exemplarily. Only results for one participant did not reject the null hypothesis during a run with normal task difficulty (*p* = .22). In 30 out of 34 runs, the lag was determined at lag = 1 indicating that HRV precedes BCI control performance by 10 s. In only four cases lags were larger with up to 70 s. However, it should be noted that 10 s was the lowest resolvable time interval as duration of one task trial (5 s) in combination with a subsequent intertrial interval (randomized between 4-6 s) was on average 10 s long. Therefore, windows were shifted by 10 s each and thus determined the respective time resolution.

**Figure 4:**
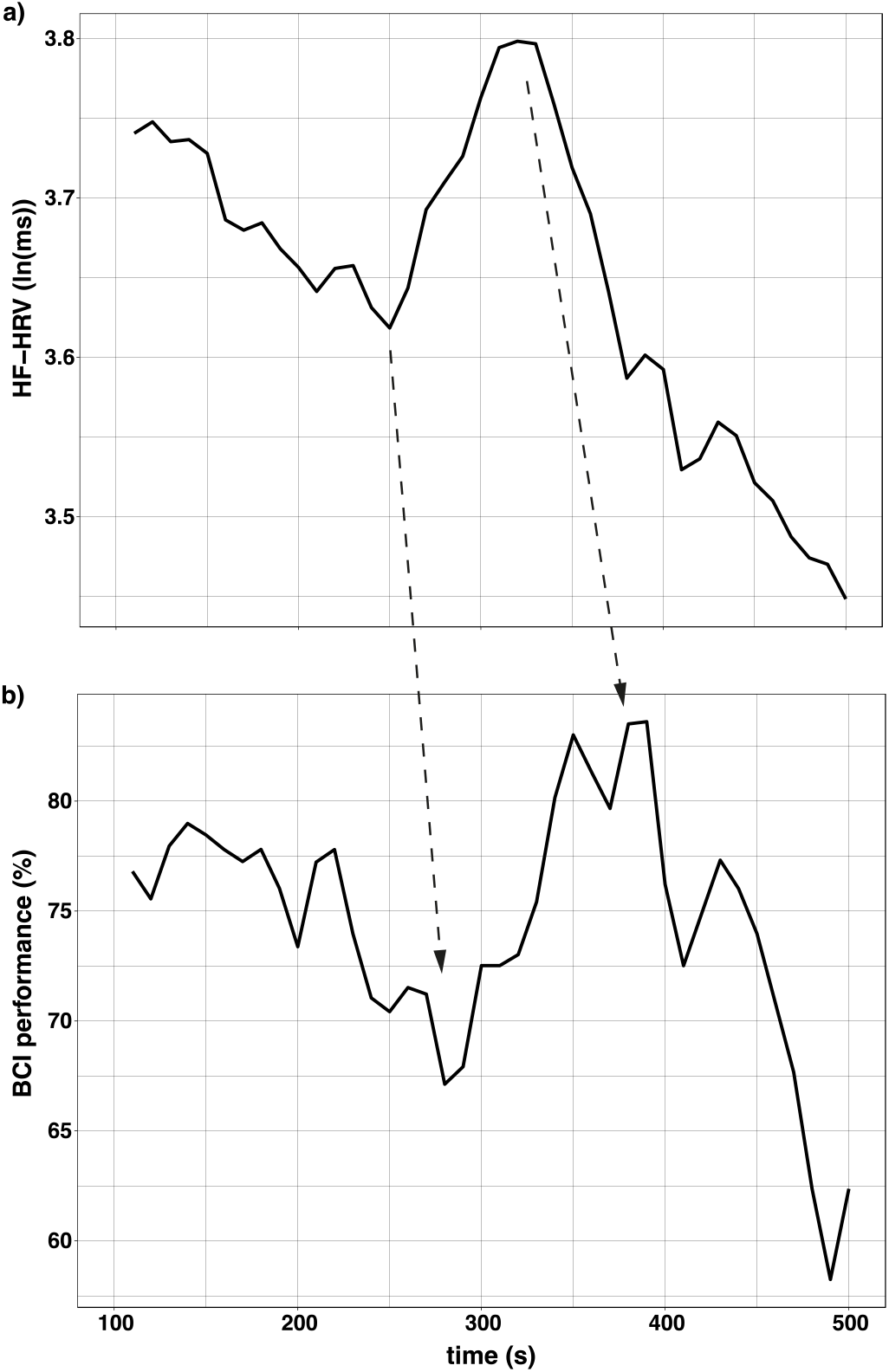
Time series of: **a)** high-frequency (HF-) heart rate variability (HRV) and **b)** brain-computer interface (BCI) performance of one participant. Dashed arrows highlight fiducial turning points, where predictive characteristics of HRV is apparent.

## 4. Discussion

Using a SMR-ERD-based BCI to control an assistive exoskeleton on a regular basis is a promising tool to trigger neural recovery in stroke neurorehabilitation. However, BCI control tasks, e.g., MI or attempting to move the paralyzed fingers, are cognitively demanding and can thus lead to inattention and fatigue, especially in stroke survivors suffering from cognitive impairments and post-stroke fatigue. As a consequence, BCI performance deteriorates over time causing frustration and reduced motivation to engage BCI control on a regular basis. Currently, there is no standardized strategy in the BCI field to anticipate decline in SMR-ERD BCI control performance online and to adapt the control paradigm accordingly. Even though numerous studies have shown that changes in HRV are closely linked to cognitive processes, it was unknown whether HRV can predict BCI control performance within a single session.

We found that during the first run with normal task difficulty, BCI performance over time remained unchanged (*p* = .176). However, there was a decline by −0.6 %/10 s (*p* = 0) during the second, more difficult run (task difficulty gradually increased, Figure 2). Since only healthy and well-performing participants were selected in a pre-screening process (see section 2.1), a decline in performance during the first run, e.g., due to fatigue, was not expected. However, the slight trend (−0.09 %/10 s) indicates that prolonged use would most likely lead to a significant decrease, too. The strong decline in the second run confirms sufficient gradual decrease of SMR-ERD detection threshold. Of course, such extreme increase in task difficulty does not reflect a normal BCI training situation, but should simulate a state of increasing exhaustion, which stroke survivors often experience during cognitive tasks due to post-stroke fatigue. Equal to the results of BCI performance, HRV was unaltered during the normal run (*p* = .157) but attenuated almost linearly within the second run (*p* < .001, Figure 3). These results demonstrate the specificity of HRV changes relative to BCI performance.

Further, causality analysis revealed that HRV *Granger-causes* BCI performance (*p* < .001) for both levels of task difficulty. The results suggest that HRV is a specific biomarker to predict decline in BCI performance.

In this context, Granger causality should not be confused with *causality* in the more general sense; that a decline in HRV directly causes a decline in BCI performance (Bunge, 2017). The result can rather be interpreted by assuming that incorporating past information of HRV substantially improves prediction of BCI performance in contrast to a prediction based on past information of BCI performance only.

These findings are in line with the neurovisceral model of Thayer et al. (2009) suggesting a close interplay between cognitive performance, HRV and prefrontal neural function that is important for executive functions including attentional control as well as motivational and emotional regulation. Moreover, the results have substantial implications on novel neuroadaptive control paradigms to individualize and optimize stroke neurorehabilitation protocols. HRV as potential control parameter for adaptation has several advantages. In contrast to EEG-based monitoring, e.g., for workload, heartbeat analysis is very simple to record and to analyze. While EEG measures usually require a time-consuming calibration to account for inter-individual variations in topographical and frequency representations of fatigue, heartbeat and for HRV analysis relevant R-peaks detection equals among all humans. To utilize this measure for individual adaptions, a baseline in HRV can easily be determined by expressing the measure as ratio score relative to task onset and define a critical linear threshold, e.g., −5 % to task onset. Hence, no time-intensive calibration is necessary. A shortcoming of this study was that ECG was recorded according to the standard lead I representation of Einthoven and is, thus, not optimal for daily home use. However, since heartbeat is one of the strongest biosignals in the human body, it is very easy to record this measure with wearables at well accessible positions, e.g., at the wrist. It is even possible to extract heart activity from the EEG, which needs to be recorded for BCI exoskeleton control, e.g., with an EEG headset.

Closed-loop neuroadaptive technology was shown to improve user performance in demanding human-computer interaction tasks. Faller et al. (2019) showed that flight duration in a boundary-avoidance task (BAT) was prolonged when dynamically adjusted EEG-based auditory neurofeedback reduced individual’s arousal state. They argued that a reduction in arousal improves performance according to the right side of the Yerkes & Dodson law stating a relation between performance and arousal describing an inverted u-shape. Importantly, the study showed that HRV increased again when arousal was downregulated by the user based on individual neurofeedback. While in our open-loop study, the opposite effect on the right side of the Yerkes & Dodson law was investigated, i.e., decrease in performance and HRV, Faller’s study indicates that HRV is suitable for closed-loop applications as HRV recovered instantaneously when arousal was reduced (Lehrer and Gevirtz, 2014, Faller et al., 2019).

Further studies are needed to investigate how to use and process the predictive HRV information. One possibility is an open-loop system to simply interrupt or stop the BCI training as soon as the stroke survivor exceeds a predefined critical HRV attenuation threshold. This would avoid frustrating and enhance rewarding experiences, e.g., to increase motivation for further training sessions. Another possibility is a closed-loop system where the BCI control paradigm is dynamically adapted. Several approaches for adaption are conceivable: Adjusting classification thresholds, recalibrating the BCI system on the fly, or applying external stimuli. It was shown, for example, that vibro-tactile feedback time-locked to heartbeat has a calming effect and might help to stabilize BCI performance (T. Azevedo et al., 2017). Nevertheless, both methods, open-loop or closed-loop, are promising to individualize BCI stroke training.

Another limitation of this study was that only the right side of the Yerkes & Dodson law was investigated. This means the behavior of HRV was only evaluated when BCI performance decreased. This was achieved by the pre-screening procedure ensuring to have a homogenous group of relatively good performers who decline in control performance over time. Contrary to this, it would be very interesting to look at the left side of the inverted u-shape, expecting that low arousal, e.g., boredom or no engagement during the task at all, also reduces performance output. Thus, a slight reduction in HRV, which was often associated with mental effort (Aasman et al., 1987), could be interpreted as task motivation and linked with a positive rewarding feedback to the user. Another possible limitation of an HRV-adaptive closed-loop system could relate to be the altered autonomic nervous system (ANS) after stroke (Dorrance and Fink, 2011). Stroke survivors show cardiac abnormalities and have in general reduced HRV (Gujjar et al., 2004). To account for such low HRV, high ECG sampling rate is suggested (Berntson et al., 1997). However, further studies are needed to evaluate the implication of such pathophysiological effects.

## 5. Conclusion

The here presented study shows that HRV is a specific biomarker to predict decline in SMR-based BCI performance. The predictive characteristics of HRV is an important prerequisite for the development of neuro-adaptive BCI control paradigms, e.g., in stroke neurorehabilitation. Optimized BCI training based on individual changes in HRV can account for patient-dependent cognitive capabilities to potentially enhance rehabilitation efficacy. For instance, frustration can be prevented as soon as decline in BCI performance is predicted. Implementation of such neuro-adaptive technology in an online BCI control paradigm is on the way.

## Notes

### Competing Interest Statement

The authors have declared no competing interest.

